# New body typology method based on the combination of coupling motion of pelvis, thorax, and cranium

**DOI:** 10.1101/2023.07.16.549262

**Authors:** Masahiro Niitsuma, Sho Imai, Takahiko Mimuro

## Abstract

The purpose of this study is to deepen the understanding of unconscious intrinsic individuality by proposing and verifying the hypothesis of “Integrated Spine Coupling” which is defined by the combination of the way of the coupling motion of 3 major units of our body: pelvis, thorax, and cranium. This paper, as the first step of the verification, examines how each of the three units — pelvis, thorax, and cranium as the representative units of cervical, thoracic, and lumber spines — produces a coupling motion during body rotational movement, and how the types of combinations are related to other movements such as breathing. The measurement results suggest that there exist only four types of coupling motion patterns and it significantly affects the way they breath and move their ankle and toe.

## Introduction

In recent years, there has been a growing emphasis on diversity in society, and the concept of Universal Design has been widely used in the design of social infrastructure that considers diversity. Universal design aims to make things as accessible as possible to a wide range of people, regardless of age, gender, ability, or circumstances [1]. It has been incorporated into various fields such as architecture, housing, public transportation, industrial products, and school education. However, universal design often categorizes people based on criteria such as age, illness, or disability, and tends to generalize the bodies of able-bodied individuals. There is also a tendency to fit limited criteria such as appearance and efficiency based on average values as the optimal solution. For example, the “correct way of walking” is often described as straightening the back, tucking in the chin, and slightly raising the gaze, but in reality, diverse bodies may not have a common solution.

The study of human body movements is conducted in several fields including: 1) kinematics - analyzing the movement of various body parts during a walking cycle through observation [2], 2) dynamics - reproducing human walking movements by generating muscle forces for each joint using joint muscle force functions and applying them to dynamics [3], and 3) functional anatomy - analyzing the functions and control of various body parts during each walking cycle [4]. Even with a single movement like walking, multidisciplinary studies are conducted, but they often focus on partial studies such as measuring the impact on specific parts of the lower limbs and tend to optimize for the average value, lacking discussion on the diversity of body movements.

Furthermore, the interplay of the entire body movement was analyzed based on the link-segment model, revealing patterns of human movement and motion. Rehabilitation applications have been developed since the late 1990s, and clinical fields have conducted analyses on the effects of abnormalities in specific body parts on other parts [5]. The notion of coupling motion has been investigated concerning the motion of the spinal chain [6]. However, experimental results concerning the coupling motion reported in the several studies, are often conflicting. In this study, in order to deepen the understanding of the diversity of human body movements, that have not been clearly elucidated in previous research, an new body typology method based on an Integrated Spinal Coupling Hypothesis (ISCH) is proposed, and a partial verification of them is conducted.

## Materials and methods

### Integrated Spinal Coupling Hypothesis

#### ISCH is consisted of the following three assumptions

1. Every body movement involves the coordination of the entire body under the restriction of the structure of the spine (movement chain).
2. There exist only the limited number of patterns in the movement chain (typification).
3. This pattern is unique to each individual and this pattern is characterized by the coupling motion of the spine (habituation).

As the movement chains of bodies influenced by the Integrated Spinal Coupling (ISC) are diverse, it would be ideal to investigate as many movement chain as possible. However, as a preliminary verification, this study first measures and verifies the existence and types of Integrated Spinal Coupling involving the cervical, thoracic, and lumbar spine, and explores their correlation with other movements such as respiratory movements.

The spine primarily undergoes an S-shaped curvature to resist gravity and is firmly connected to various regions such as the pelvis, thorax, and cranium. Therefore, ISCH suggests that the directionality of the movements in the regions of the spine, especially the pelvis, thorax, and cranium, significantly affect other movements as the result of the movement chain. ISCH also suggests that this directionality is habitually unique to the individual.

In this study, to propose a new body typology method on the basis of ISCH, we focus on the directionality of the three units: pelvis, thorax, and cranium (as the representative units of cervical, thoracic, and lumber spines). A body typology method is proposed based on the response of these units to rotational movements. When the three units perform rotational movements, each unit exhibits lateral flexion and flexion/extension (often referred to as the coupling motion). Furthermore, the coupling motion can be classified into either ipsilateral (i) or contralateral rotation (c). According to this definition, there exist eight possible combinations of pelvic, thorax, and cranium units: ccc, cci, cic, cii, icc, ici, iic, iii. Therefore, it means that individuals unconsciously select one combination from these options, and this selection becomes habitual. Similar ideas to ISCH have already been implicitly practiced and proven to be effective to some extent, in practical settings. However, quantitative data has not been conducted. By explicitly verifying the body typology method based on ISCH, various studies that lead to true universal design tailored to individual body characteristics, could be conducted in the future.

Participants were recruited from September to October 2022. Authors had no access to information that could identify individual participants during or after data collection.

### Body Rotational Movements

The measurement of body rotational movements involved 22 consenting adult participants (16 males and 6 females) who met the following criteria: (1) able to perform body rotational movements without pain and independently, and (2) capable of walking independently. The participants performed the twisting movement of the body multiple times while in an upright position, and the movements were simultaneously recorded by two cameras positioned in the front and side views. During the measurement of twisting movements, the spine was divided into three units: lumbar vertebrae + sacrum + coccyx and pelvis, thoracic vertebrae + ribcage, and cervical vertebrae + skull. It was determined through camera footage and interviews whether each unit exhibited flexion/extension and lateral flexion coupling motion and, if so, whether it was ipsilateral or contralateral rotation. Fig. 1 illustrates the experiment where body rotational movements are measured, from the frontal view.

**Fig 1.**
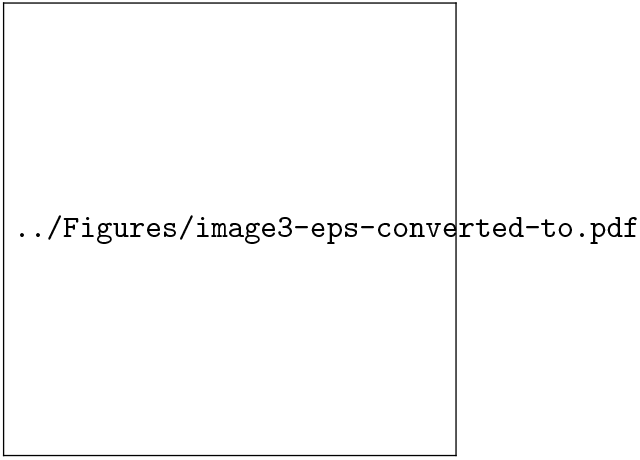
How body rotational movements are measured in the experiment.

### Respiratory movements

The measurement of respiratory movements was conducted on the participants who underwent the measurement of body rotational movements. To ensure that the participants maintained their habitual breathing pattern during the measurement, they were informed in advance that their usual breathing would be observed and that there was no right or wrong way to breathe. This prevented the participants from consciously altering their breathing patterns. During the measurement, the participants were instructed to relax their whole body and perform slow, deep inhalation and exhalation to their maximum capacity, maintaining the same posture and relaxation. While keeping their hands lightly placed on their abdomen (below the thorax), they were asked to perform slow inhalation and exhalation in the same manner. During the initial breath (shallow breath), the participants were questioned about whether their abdomen contracted or expanded. The process of measuring respiratory movements is illustrated in Fig. 2.

**Fig 2.**
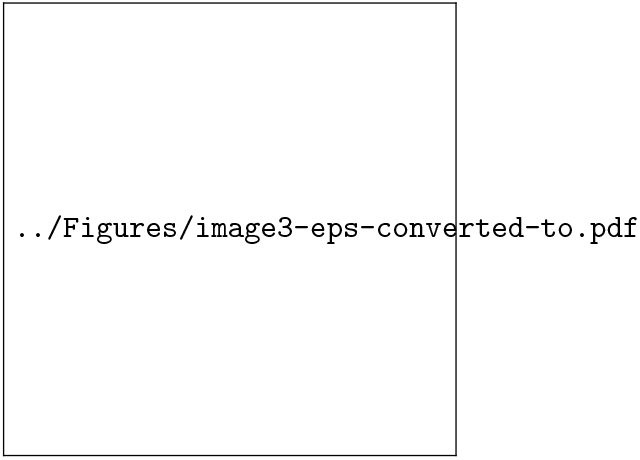
How respiratory movements are measured in the experiment.

### Ankle and toe movements

The measurement of toe and ankle movements was conducted on subjects who have undergone body rotation measurement, following the procedure outlined below:

1. In a seated position, observe the sequential state of the toes while performing ankle dorsiflexion and plantar flexion.
2. Conduct an interview to assess the ease of performing the movements and habitual patterns.
3. Raise one leg and observe the sequential state of the ankle while performing toe dorsiflexion and plantar flexion.
4. Raise one leg and observe the sequential state of the ankle while performing toe dorsiflexion and plantar flexion.

## Results

### Body Rotational Movements

Table 1 shows the results of the measurements, confirming the presence of ISC in all 22 participants. Although anatomical confirmation of the spine was not possible in this study, visual observation and interviews with the participants indicated that during body rotation, significant flexion and extension (coupling motions) were observed in the major regions of the cranium, thorax, and pelvis. This suggests that the individual parts of the spine, which are strongly connected to these three regions, also exhibit similar coupling motions. Additionally, the participants were asked to experience both ipsilateral and contralateral coupling motions during the rotation of the pelvis, thorax, and cranium, and they were questioned about the differences in each of movement compared to their habitual actions. The results showed that the ease of movement varied significantly among the participants, and many of them experienced a notable discomfort when using unfamiliar direction. These findings suggest that coupling motion movements may be habitual and involuntary strategies. Furthermore, it was found that the combination of coupling motions in the three units (pelvis, thorax, and cranium) can be only following five types:, cci, cic, icc, ici, iii, indicating the coupling motion of the cranium unit is determined by the combination of the coupling motion of pelvis and thorax units.

**Table 1.**
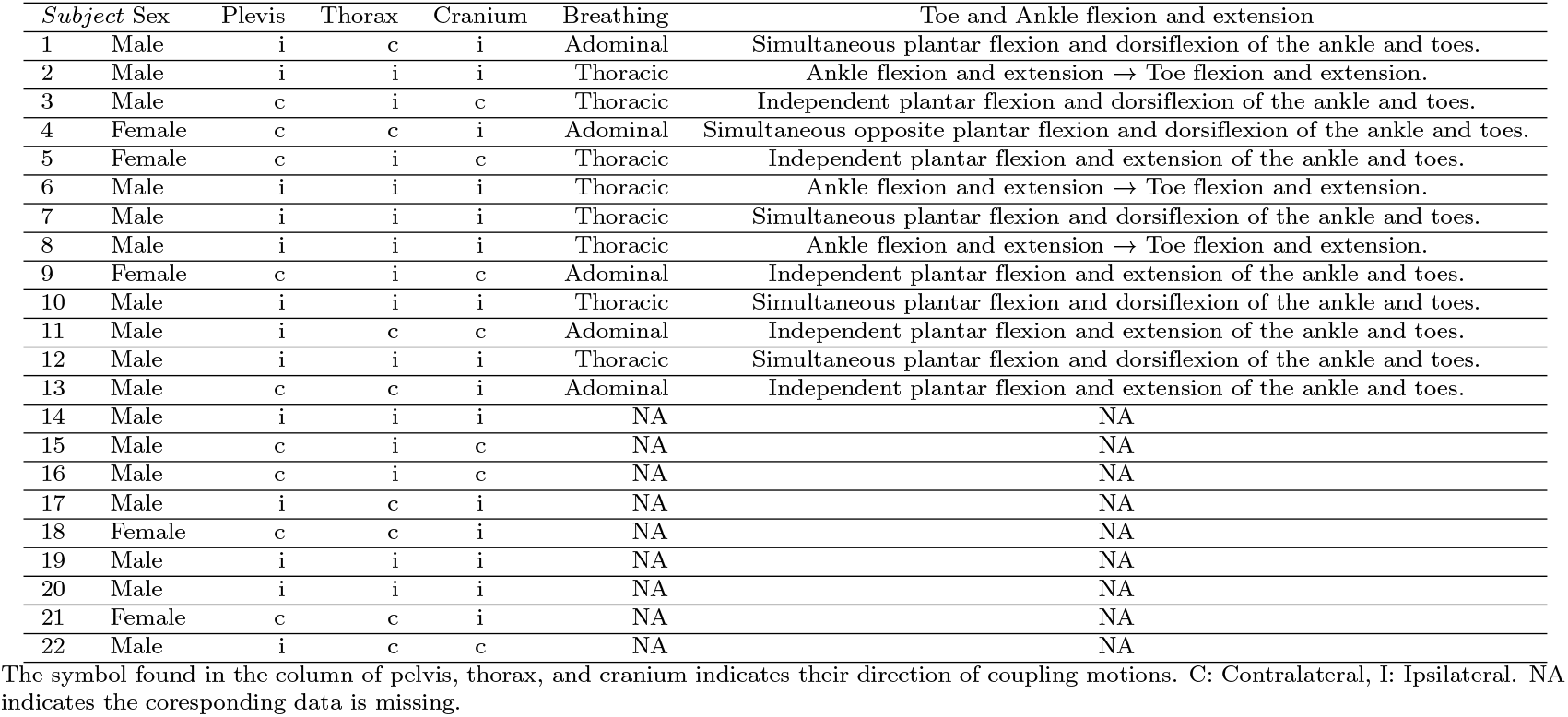
Combination of the type of coupling motions and its relation to other motions.

### Respiratory movements

The results of the measurement of the participants’ breathing patterns are shown in Table 1. Among the four participants who exhibited contralateral rotation of the thorax unit, all four (100%) performed diaphragmatic inhalation followed by chest inhalation, and exhaled in the sequence of chest exhalation followed by diaphragmatic exhalation. On the other hand, among the participants with ipsilateral rotation of the thorax unit during body rotation, eight out of nine participants (89%) performed chest inhalation followed by diaphragmatic inhalation, and exhaled in the sequence of diaphragmatic exhalation followed by chest exhalation. The remaining one participant (11%) performed diaphragmatic inhalation followed by chest inhalation and exhaled in the sequence of chest exhalation followed by diaphragmatic exhalation. Questionnaires were conducted to confirm habitual breathing patterns and all participants reported having habitual breathing patterns. Additionally, among the participants who exhibited ipsilateral rotation of the thorax unit and performed diaphragmatic inhalation followed by chest inhalation, one participant mentioned having experience playing wind instruments and receiving instruction on diaphragmatic breathing in the past.

### Ankle and toe movements

The measurement results of ankle and toe movements in the subjects are presented in Table 1. Among the four types classified based on the ISCH, the cc (Pelvis: Contralateral, thorax: Contralateral) type exhibited reverse sequential movement of ankle dorsiflexion and plantar flexion in 50% of the subjects, while the other 50% showed non-sequential movement. The ci (Pelvis: Contralateral, Thorax: Ipsilateral) type showed non-sequential ankle dorsiflexion and plantar flexion in 100% of the subjects. The II (Pelvis: Ipsilateral, Thorax: Ipsilateral) type demonstrated sequential movement of ankle dorsiflexion and plantar flexion in the same direction in 100% of the subjects. The ic (Pelvis: Ipsilateral, Thorax: Contralateral) type exhibited sequential ankle dorsiflexion and plantar flexion in the same direction in 50% of the subjects, while the remaining 50% showed non-sequential movement. Interviews regarding the movements confirmed that all sequential/non-sequential patterns were habitual and involuntary.

## Discussion

### Body Rotational Movements

According to the measurement results, the presence of ISC was confirmed in all 22 participants. Although anatomical confirmation of the spine was not possible in this study and confirmation was based on visual observation and participant interviews, it was suggested that significant flexion (coupling motion) in the anterior-posterior and lateral directions was observed during body rotation in at least the major regions of the cranium, thorax, and pelvis. It was also suggested that each segment of the spine, which is strongly interconnected with these three regions, undergoes similar coupling motions. Furthermore, participants were asked to experience both ipsilateral and contralateral coupling motions during the rotation of the pelvis, thorax, and cranium, and were questioned about the ease of movement and differences compared to their usual movements. The results revealed clear differences in the ease of movement among participants, with many participants expressing significant discomfort when using unfamiliar directionality. These findings are consistent with ISCH concerning typification and habituation. Moreover, it was found that the coupling motion of the cranium unit is determined by the combination of the coupling motion of pelvis and thorax units.

### Respiratory movements

Based on the measurement results, it can be inferred that the breathing pattern is highly influenced by the coupling motion of the thoracic unit. In this study, during the initial phase of inhalation, all participants with contralateral rotation of the thoracic unit performed diaphragmatic breathing, while most participants with ipsilateral rotation (except for one) performed thoracic breathing during the initial phase of inhalation. Upon interviewing the exceptional participant, it was revealed that they had received instruction and practice in diaphragmatic breathing as part of their training in playing wind instruments. In the context of the ISC, the rules of the ISCH are considered to be applicable to the initial stage of human movement and are expected to undergo changes due to various external factors in real-life situations.

### Ankle and toe movements

From the measurement results, it can be inferred that there is a strong correlation between the coupling motion strategy of the ISC and the movement of ankle and toe dorsiflexion/plantar flexion. Among the subjects with ipsilateral pelvic rotation, all subjects with ipsilateral thoracic rotation and half of the subjects with contralateral thoracic rotation exhibited ankle and toe dorsiflexion/plantar flexion in the same sequential direction (the remaining half showed non-sequential movement). This suggests a high possibility that in the coupling motion strategy of ipsilateral pelvic rotation in the ISC, muscles spanning two joints such as the long toe flexors and long toe extensors play a dominant role in producing dorsiflexion/plantar flexion. On the other hand, regarding the strategy of contralateral pelvic rotation, half of the subjects with ipsilateral thoracic rotation and half of the subjects with contralateral thoracic rotation showed non-sequential movement, while the remaining half exhibited reverse sequential movement. These results support the notion in the ISC that in the strategy of contralateral pelvic rotation, single-joint muscles predominantly induce independent or reverse sequential responses in ankle and toe dorsiflexion/plantar flexion.

## Conclusion

The verification of the ISCH involved classifying body rotational movements and then measuring respiratory movements, ankle and toe movements to demonstrate the correlation between spinal coupling patterns and habitual movement strategies. The results suggested a significant correlation between spinal coupling patterns and respiratory movements as well as ankle and toe movements. Through the analysis of the measurement results in this study, the verification of the ISCH was partially achieved. This provides significant insights into the typification of bodily movements, which has been a crucial challenge that has remained unclear until now. This knowledge holds relevance in various fields related to the human body, including design considerations encompassing the body and its environment. The verification of ISCH highlights the significance of acknowledging habitual movement strategies influenced by individual differences in spinal structure, challenging the conventional point of views based on the generalization of bodily movements.

To improve the verification of ISCH by overcoming the limitations of this study, there are the following four aspects to be addressed. First, the current experiment involved a relatively small sample size of 22 participants, which may have limited the detailed analysis, particularly for specific ISC patterns with fewer participants. Additional measurements need to be conducted to address this limitation. Second, this study focused on measuring respiratory movements, ankle and toe movements, to examine the relationship between ISC patterns and habitual movement strategies. However, it is also necessary to investigate other habitual movements that primarily involve the upper extremities, and the actions involving the use of tools. Further research is needed to quantitatively evaluate the correlation between these various movements and ISP patterns. Third, systems approach would be applied to create models that enables deductive interpretations of the result obtained by the experiment, thereby possibly integrating several pieces of knowledge concerning our body structure and movements, which have previously been studied in a fragmented manner, as shown in the Introduction. Fourth, it is necessary to conduct measurements using not only visual observation, but also force plates or other quantitative measurement instruments to enable more objective analysis and description.

---

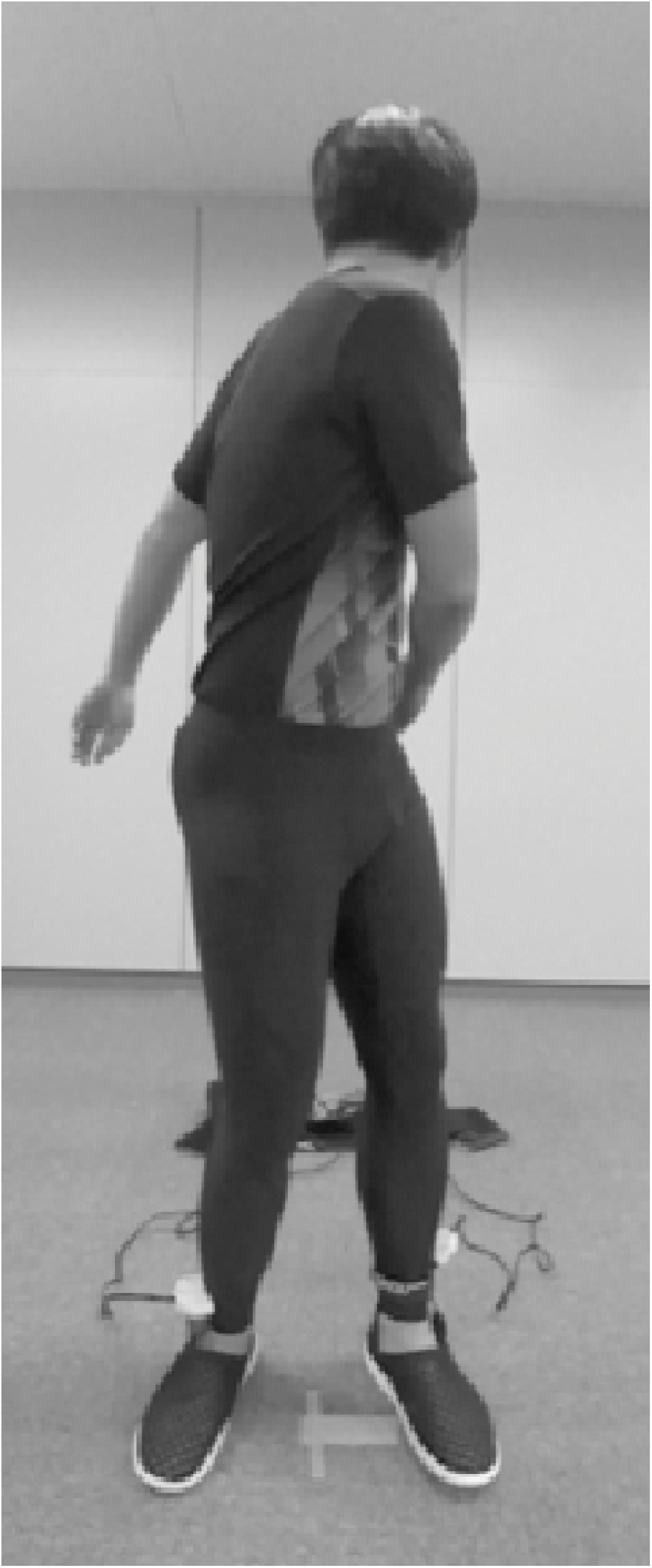

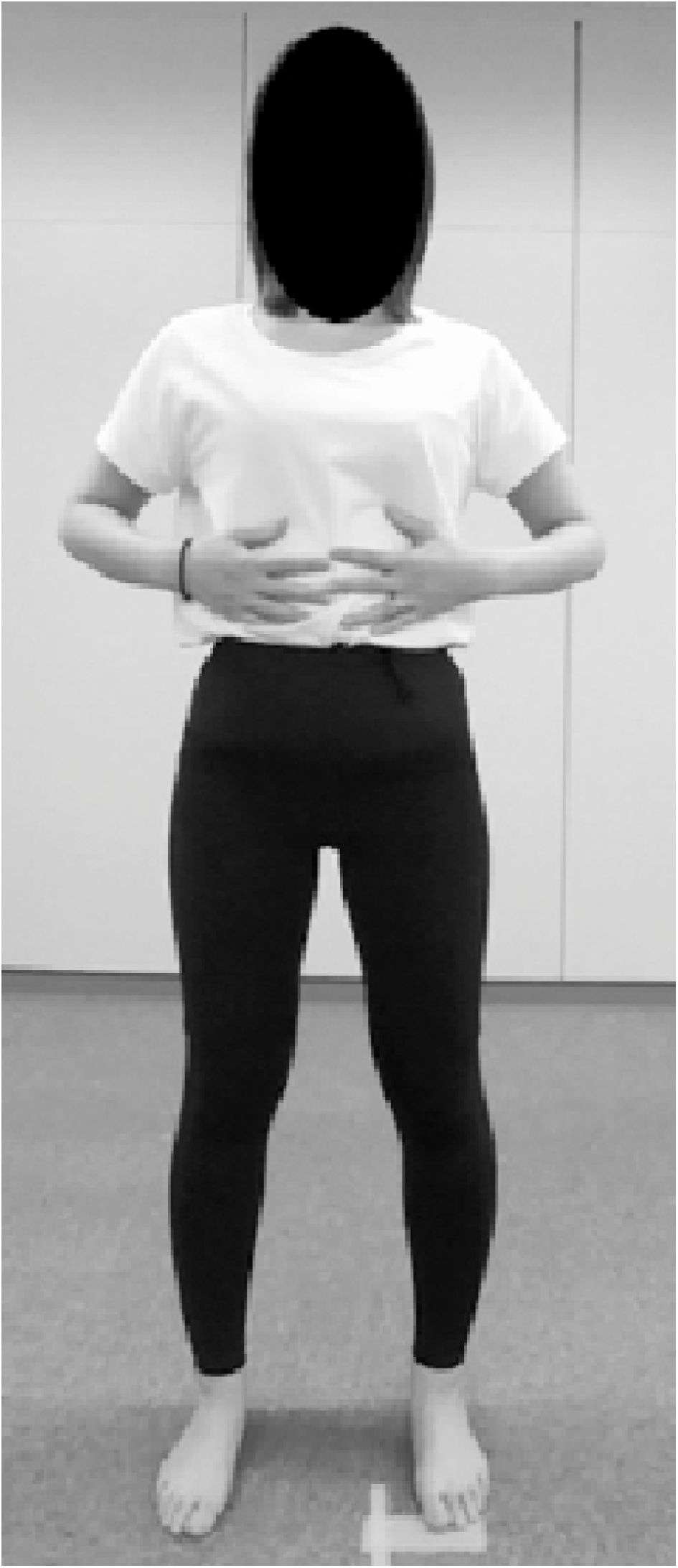

## Notes

### Competing Interest Statement

The authors have declared no competing interest.

